# Multivariate Genomewide Association Analysis by Iterative Hard Thresholding

**DOI:** 10.1101/2021.08.04.455145

**Authors:** Benjamin B. Chu, Seyoon Ko, Jin J. Zhou, Aubrey Jensen, Hua Zhou, Janet S. Sinsheimer, Kenneth Lange

**Affiliations:** Department of Computational Medicine, David Geffen School of Medicine at UCLA, Los Angeles, USA; Department of Biostatistics, Fielding School of Public Health at UCLA, Los Angeles, USA; Division of Epidemiology and Biostatistics, University of Arizona, USA; Department of Human Genetics, David Geffen School of Medicine at UCLA, Los Angeles, USA; Department of Statistics at UCLA, Los Angeles, USA

**Keywords:** sparsity, Gaussian, iterative hard thresholding, Julia

## Abstract

In genome-wide association studies (GWAS), analyzing multiple correlated traits is potentially superior to conducting multiple univariate analyses. Standard methods for multivariate GWAS operate marker-by-marker and are computationally intensive. We present a penalized regression algorithm for multivariate GWAS based on iterative hard thresholding (IHT) and implement it in a convenient Julia package `MendelIHT.jl` (https://github.com/OpenMendel/MendelIHT.jl). In simulation studies with up to 100 traits, IHT exhibits similar true positive rates, smaller false positive rates, and faster execution times than `GEMMA`’s linear mixed models and `mv-PLINK`’s canonical correlation analysis. On UK Biobank data, our IHT software completed a 3-trait joint analysis in 20 hours and an 18-trait joint analysis in 53 hours, requiring up to 80GB of computer memory. In short, our software enables geneticists to fit a single regression model that simultaneously considers the effect of all SNPs and dozens of traits.

## 2 Introduction

Current statistical methods for genome-wide association studies (GWAS) can be broadly categorized as single variant or multi-variant in their genomic predictors. Multi-variant sparse models ignore polygenic background and assume that only a small number of single-nucleotide polymorphisms (SNPs) are truly causal for a given trait. Model fitting is typically accomplished via regression with penalties such as the least absolute shrinkage and selection operator (LASSO) [3, 34, 42, 48, 46], minimax concave penalty (MCP) [7, 45], iterative hard thresholding (IHT) [8, 18], or Bayesian analogues [15]. Linear mixed models (LMM) dominate the single variant space. LMMs control for polygenic background while focusing on the effect of a single SNP. LMMs are implemented in the contemporary programs GEMMA [51], BOLT [25], GCTA [17, 43], and SAIGE [50]. The virtues of the various methods vary depending on the genetic architecture of a trait, and no method is judged uniformly superior [13].

Although there is no consensus on the best modeling framework for single-trait GWAS, there is considerable support for analyzing multiple correlated traits jointly rather than separately [13, 31, 41]. When practical, joint analysis (a) incorporates extra information on cross-trait covariances, (b) distinguishes between pleiotropic and independent SNPs, (c) reduces the burden of multiple testing, and (d) ultimately increases statistical power. Surprisingly, simulation studies suggest these advantages hold even if only one of multiple traits is associated with a SNP or if the correlation among traits is weak [13]. These advantages motivate the current paper and our search for an efficient method for analyzing multivariate traits.

Existing methods for multivariate-trait GWAS build on the polygenic model or treat SNPs one by one. For instance, GEMMA [52] implements multivariate linear mixed models (mvLMM), mv-PLINK [11] implements canonical correlation analysis, and MultiPhen [30] and Scopa [27] invert regression so that the genotypes at a single SNP become the trait and observed traits become predictors. Due to their single-variant nature, these methods cannot distinguish whether a SNP exhibits a true effect on the trait or a secondary association mediated by linkage disequilibrium. As a result, many correlated SNPs near the causal one are also selected. This inflates the false positive rate unless one applies fine-mapping strategies [37] in downstream analysis to distill the true signal. Joint regression methods like IHT and LASSO are less susceptible to finding SNPs with only secondary association because all SNPs are considered simultaneously.

To our knowledge, there are no sparse regression methods for multivariate-trait GWAS. In this paper, we extend IHT [6] to the multivariate setting and implement it in the Julia [5] package `MendelIHT.jl`, part of the larger OpenMendel statistical genetics ecosystem [49]. We have previously demonstrated the virtues of IHT compared to LASSO regression and single-SNP analysis for univariate GWAS [8, 18]. Since IHT assumes sparsity and focuses on mean effects, it is ill suited to capture polygenic background as represented in classic variance components models. In the sequel we first describe our generalization of IHT. Then we study the performance of IHT on simulated traits given real genotypes. These simulations explore the impact of varying the sparsity level *k* and the number of traits *r*. To demonstrate the potential of IHT on real large-scale genomic data, we also apply it to 3 hypertension related traits and 18 metabolomic traits from the UK Biobank. These studies showcase IHT’s speed, low false positive rate, and scalability to large numbers of traits. Our concluding discussion summarizes our main findings, limitations of IHT, and questions worthy of future research.

## 3 Materials and Methods

### 3.1 Model Development

Consider multivariate linear regression with *r* traits under a Gaussian model. Up to a constant, the loglikelihood 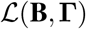 for *n* independent samples is

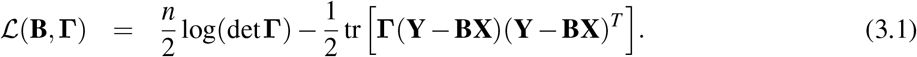

The loglikelihood 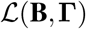 is a function of the *r* × *p* regression coefficients matrix **B** and the *r* × *r* unstructured precision (inverse covariance) matrix **Γ**. Furthermore, **Y** is the *r* × *n* matrix of traits (responses), and **X** is the *p* × *n* design matrix (genotypes plus non-genetic predictors). All predictors are treated as fixed effects.

IHT maximizes 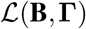 subject to the constraints that *k* or fewer entries of **B** are non-zero and that **Γ** is symmetric and positive definite. Optimizing 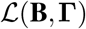 with respect to **B** for **Γ** fixed relies on three core ideas. The first is gradient ascent. Elementary calculus tells us that the gradient 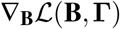 is the direction of steepest ascent of 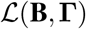 at **B** for **Γ** fixed. IHT updates **B** in the steepest ascent direction by the formula 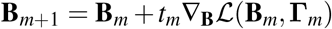, where *t*_*m*_ > 0 is an optimally chosen step length and (**B**_*m*_, **Γ**_*m*_) is the current value of the pair (**B**, **Γ**). The gradient is derived in the appendix as the matrix

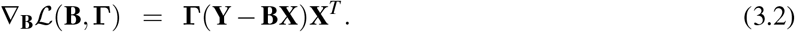

The second core idea dictates how to choose the step length *t*_*m*_. This is accomplished by expanding the function 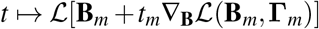 in a second-order Taylor series around (**B**_*m*_, **Γ**_*m*_). Our appendix shows that the optimal *t*_*m*_ for this quadratic approximant is

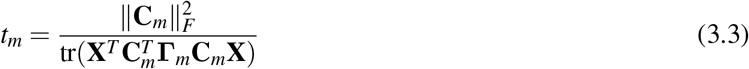

given the abbreviation 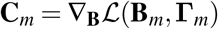. The third core idea of IHT involves projecting the steepest ascent update 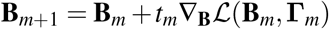 to the sparsity set *S*_*k*_ = {B: ║B║_0_ ≤ *k*}. The projection operator 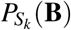 sets to zero all but the largest *k* entries in magnitude of **B**. This goal can be achieved efficiently by a partial sort on the vectorized version vec(**B**_*m*+1_) of **B**_*m*+1_. For all predictors to be treated symmetrically in projection, they should be standardized to have mean 0 and variance 1. Likewise, in cross-validation of *k* with mean square error prediction, it is a good idea to standardize all traits.

To update the precision matrix **Γ** for **B** fixed, we take advantage of the gradient

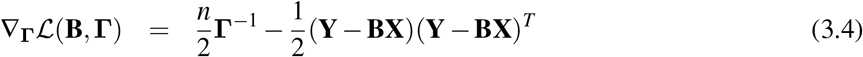

spelt out in the appendix. At a stationary point where 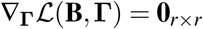, the optimal **Γ** is

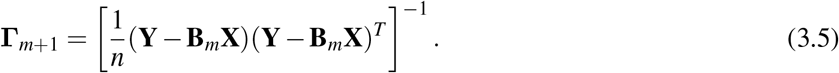

Equation (3.5) preserves the symmetry and positive semidefiniteness of **Γ**_*m*_. The required matrix inversion is straightforward unless the number of traits *r* is exceptionally large. Our experiments suggest solving for **Γ**_*m*+1_ exactly is superior to running full IHT jointly on both **B** and **Γ**. Figure 1 displays our block ascent algorithm.

**Figure 1:**
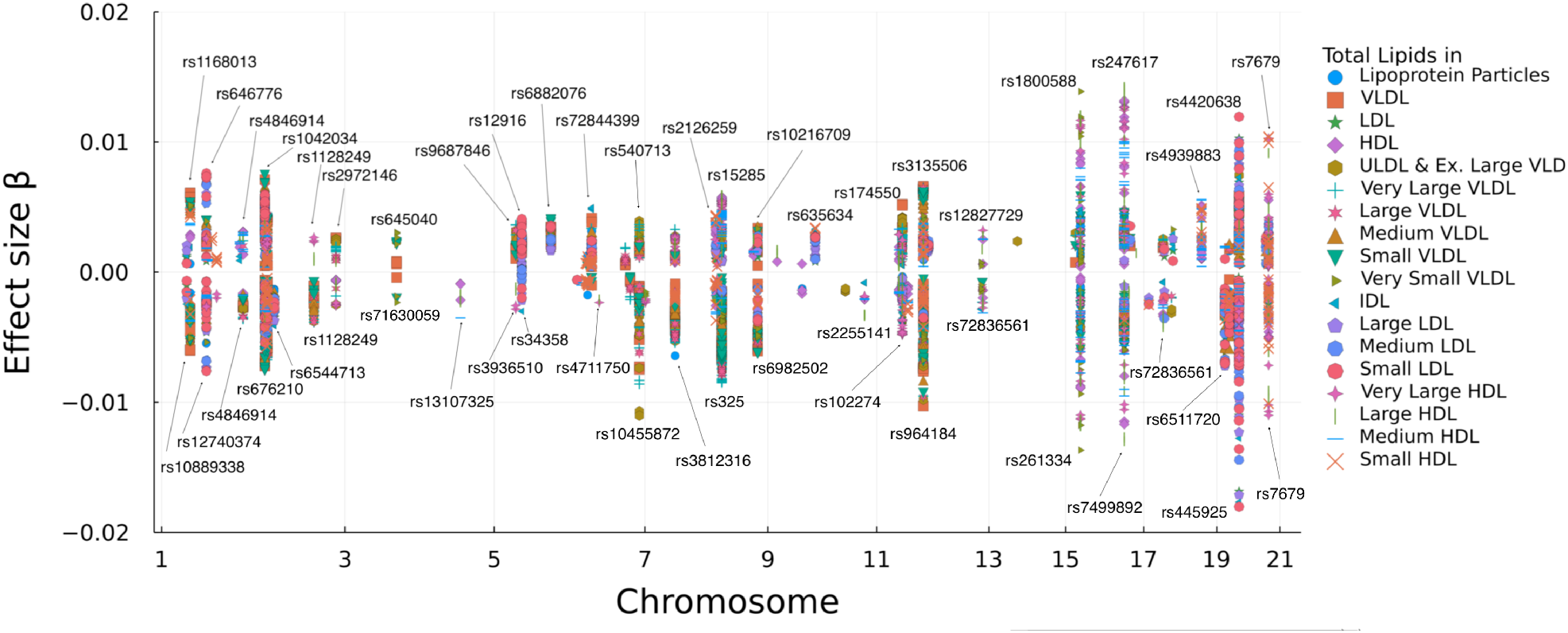
An 18-trait joint analysis on UK Biobank’s metabolomic traits using multivariate IHT. The effect size for each trait is plotted against its chromosome position. The larger effect sizes are labeled with their SNP names. Note one unit increase in effect size does not directly translate to one unit increase of lipids levels in its original scale because all traits were log-transformed and standardized.

#### Algorithm 1

Block Ascent Multivariate Iterative Hard-Thresholding

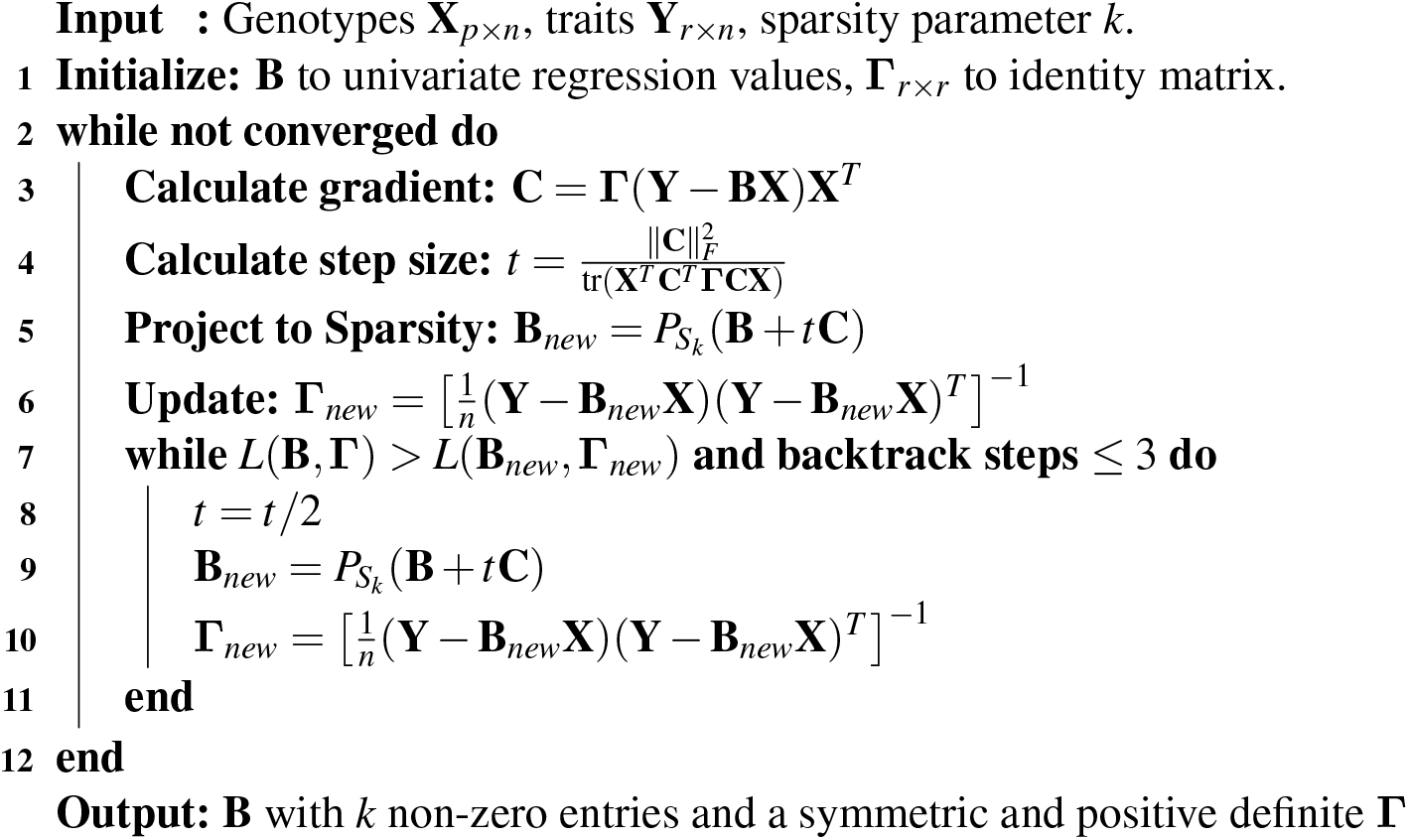

### 3.2 Linear Algebra with Compressed Genotype Matrices

We previously described how to manipulate PLINK files using the `OpenMendel` module `SnpArrays.jl` [49], which supports linear algebra on compressed genotype matrices [8]. We now outline several enhancements to our compressed linear algebra routines.

#### Compact genotype storage and fast reading

A binary PLINK genotype [33] stores each SNP genotype in two bits. Thus, an *n* × *p* genotype matrix requires 2*np* bits of memory. For bit-level storage `Julia` [5] supports the 8-bit unsigned integer type (`UInt8`) that can represent four sample genotypes simultaneously in a single 8-bit integer. Extracting sample genotypes can be achieved via bitshift and bitwise and operations. Genotypes are stored in little endian fashion, with 0, 1, 2, and missing genotypes mapped to the bit patterns 00, 10, 11, and 01, respectively. For instance, if a locus has four sample genotypes 1, 0, 2, and missing, then the corresponding `UInt8` integer is 01110010 in binary representation. Finally, because the genotype matrix is memory-mapped, opening a genotype file and accessing data are fast even for very large files.

#### SIMD-vectorized and tiled linear algebra

In IHT the most computationally intensive operations are the matrix-vector and matrix-matrix multiplications required in computing gradients. To speed up these operations, we employ SIMD (single instruction, multiple data) vectorization and tiling. On machines with SIMD support such as AVX (Advanced Vector Extensions), our linear algebra routine on compressed genotypes is usually twice as fast as BLAS 2 (Basic Linear Algebra Subroutines) [24] calls with an uncompressed numeric matrix and comparable in speed to BLAS 3 calls if **B** is tall and thin.

Computation of the matrix product **C** = **AB** requires special care when **A** is the binary PLINK-formatted genotype matrix and **B** and **C** are numeric matrices. The idea is to partition these three matrices into small blocks and exploit the representation **C**_*ij*_ = Σ_*k*_ **A**_*ik*_**B**_*kj*_ by computing each tiled product **A**_*ik*_**B**_*kj*_ in parallel. Because entries of a small matrix block are closer together in memory, this strategy improves cache efficiency. The triple `for` loops needed for computing products **A**_*ik*_**B**_*kj*_ are accelerated by invoking Julia’s `LoopVectorization.jl` package, which performs automatic vectorization on machines with SIMD support. Furthermore, this routine can be parallelized because individual blocks can be multiplied and added independently. Because multi-threading in Julia is composable, these parallel operations can be safely nested inside other multi-threading Julia functions such as IHT’s cross-validation routine.

### 3.3 Simulated Data Experiments

Our simulation studies are based on the chromosome 1 genotype data of the Northern Finland Birth Cohort (NFBC) [35]. The original NFBC1966 data contain 5402 samples and 364,590 SNPs; 26,906 of the SNPs reside on chromosome 1. After filtering for samples with at least 98% genotype success rate and SNPs with missing data less than 2%, we ended with 5340 samples and 24,523 SNPs on chromosome 1. For *r* traits, traits are simulated according to the matrix normal distribution [9, 12, 44] as

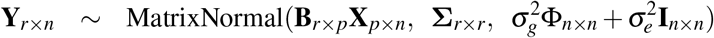

using the `OpenMendel` module `TraitSimulation.jl` [16]. Here **X** is the chromosome 1 NFBC *p* × *n* genotype matrix with *n* samples aligned along its columns. The matrix **B** contains the true regression coefficients *b*_ij_ uniformly drawn from {0.05, 0.1, …, 0.5} and randomly set to 0 so that *k*_*true*_ entries *b*_*ij*_ survive. In standard mathematical notation, ║**B**║_0_ = *k*_*true*_. Note the effect-size set {0.05, 0.1,…, 0.5} is comparable to previous studies [8]. To capture pleiotropic effects, *k*_*plei*_ SNPs are randomly chosen to impact 2 traits. The remaining *k*_*indep*_ causal SNPs affect only one trait. Thus, *k*_*true*_ = 2*k*_*plei*_ + *k*_*indep*_. Note it is possible for 2 traits to share 0 pleiotropic SNPs. The row (trait) covariance matrix Σ is simulated so that its maximum condition number does not exceed 10. The column (sample) covariance matrix equals 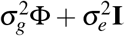, where **Φ** is the centered genetic relationship matrix (GRM) estimated by GEMMA [52]. We let 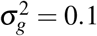 and 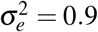, so polygenic heritability is 10%. Different combinations of *r*, *k*_*true*_, *k*_*indep*_, and *k*_*plei*_ are summarized in Table 1. Each combination is replicated 100 times. It is worth emphasizing that this generative model should favor LMM analysis.

**Table 1:** Comparison of multivariate IHT (`mIHT`) and multiple univariate IHT (`uIHT`) implemented in `MendelIHT`, canonical correlation analysis (CAA) implemented in `mv-PLINK`, and multivariate linear mixed models (mvLMM) implemented in `GEMMA` on chromosome 1 of the NFBC1966 data with simulated traits. Plei TP is the proportion of true positives for pleiotopic SNPs, Indep TP is proportion of true positives for independent SNPs, and FDR is the false discovery rate (number of false positives divided by total number selected). indicates standard deviations. *k*_*true*_ is the total number of non-zero entries in **B**, *k*_*indep*_ is the number of independent SNPs affecting only one trait, and *k*_*plei*_ is the number of pleiotropic SNPs affecting two traits. These numbers satisfy *k*_*true*_ = 2 × *k*_*plei*_ + *k*_*indep*_. Each simulation relied on 100 replicates. NA: >24 hours. (*) Only two replicates contribute to timing.

|  | Time (sec) | Plei TP | Indep TP | FDR |
| --- | --- | --- | --- | --- |
| **Set 1:** (2 traits, $k_{true} = 10$, $k_{indep} = 4$, $k_{plei} = 3$) | | | | |
| mIHT | $164.6 \pm 69.3$ | $0.92 \pm 0.16$ | $0.76 \pm 0.2$ | $0.25 \pm 0.24$ |
| uIHT | $114.9 \pm 48.6$ | $0.93 \pm 0.16$ | $0.72 \pm 0.2$ | $0.14 \pm 0.16$ |
| CCA | $152.6 \pm 57.3$ | $0.96 \pm 0.14$ | $0.78 \pm 0.2$ | $0.91 \pm 0.06$ |
| mvLMM | $307.7 \pm 121.4$ | $0.95 \pm 0.15$ | $0.76 \pm 0.2$ | $0.86 \pm 0.06$ |
| **Set 2:** (3 traits, $k_{true} = 20$, $k_{indep} = 10$, $k_{plei} = 5$) | | | | |
| mIHT | $214.4 \pm 100.1$ | $0.91 \pm 0.12$ | $0.75 \pm 0.14$ | $0.27 \pm 0.18$ |
| uIHT | $169.6 \pm 81.9$ | $0.86 \pm 0.16$ | $0.72 \pm 0.16$ | $0.16 \pm 0.12$ |
| CCA | $226.8 \pm 101.9$ | $0.95 \pm 0.09$ | $0.79 \pm 0.15$ | $0.89 \pm 0.04$ |
| mvLMM | $449.9 \pm 221.7$ | $0.93 \pm 0.1$ | $0.75 \pm 0.16$ | $0.83 \pm 0.04$ |
| **Set 3:** (5 traits, $k_{true} = 30$, $k_{indep} = 16$, $k_{plei} = 7$) | | | | |
| mIHT | $227.9 \pm 41.1$ | $0.93 \pm 0.09$ | $0.73 \pm 0.12$ | $0.22 \pm 0.12$ |
| uIHT | $213.8 \pm 45.7$ | $0.9 \pm 0.11$ | $0.69 \pm 0.12$ | $0.14 \pm 0.11$ |
| CCA | $371.5 \pm 34.0$ | $0.96 \pm 0.07$ | $0.75 \pm 0.11$ | $0.9 \pm 0.03$ |
| mvLMM | $1135.3 \pm 125.5$ | $0.94 \pm 0.09$ | $0.71 \pm 0.11$ | $0.83 \pm 0.03$ |
| **Set 4:** (10 traits, $k_{true} = 10$, $k_{indep} = 4$, $k_{plei} = 3$) | | | | |
| mIHT | $278.8 \pm 53.0$ | $0.97 \pm 0.09$ | $0.74 \pm 0.2$ | $0.23 \pm 0.17$ |
| uIHT | $245.8 \pm 34.8$ | $0.96 \pm 0.11$ | $0.7 \pm 0.23$ | $0.26 \pm 0.18$ |
| CCA | $985.1 \pm 97.5$ | $0.99 \pm 0.06$ | $0.78 \pm 0.2$ | $0.9 \pm 0.05$ |
| mvLMM | $8067.4 \pm 3900.8$ | $0.99 \pm 0.06$ | $0.74 \pm 0.18$ | $0.86 \pm 0.05$ |
| **Set 5:** (50 traits, $k_{true} = 20$, $k_{indep} = 10$, $k_{plei} = 5$) | | | | |
| mIHT | $1892.2 \pm 419.0$ | $0.93 \pm 0.12$ | $0.75 \pm 0.14$ | $0.18 \pm 0.11$ |
| uIHT | $1336.9 \pm 310.2$ | $0.92 \pm 0.11$ | $0.72 \pm 0.12$ | $0.36 \pm 0.14$ |
| CCA | $26589.1 \pm 907.7(*)$ | NA | NA | NA |
| mvLMM | NA | NA | NA | NA |
| **Set 6:** (100 traits, $k_{true} = 30$, $k_{indep} = 16$, $k_{plei} = 7$) | | | | |
| mIHT | $3699.3 \pm 410.4$ | $0.91 \pm 0.11$ | $0.71 \pm 0.11$ | $0.13 \pm 0.08$ |
| uIHT | $2353.8 \pm 212.3$ | $0.92 \pm 0.11$ | $0.7 \pm 0.1$ | $0.36 \pm 0.1$ |
| CCA | NA | NA | NA | NA |
| mvLMM | NA | NA | NA | NA |

### 3.4 Method Comparisons

In our simulation experiments, we compared `MendelIHT.jl` to `mv-PLINK` [11] and `GEMMA` [52]. The linear mixed model software `GEMMA` enjoys wide usage in genetic epidemiology. The software `mv-PLINK` is chosen for its speed. A recent review [13] rates `mv-PLINK` as the second fastest of the competing programs. The fastest method, MV-BIMBAM [38], is an older method published by the authors of `GEMMA`, so it is not featured in this study.

In simulated data experiments, all programs were run within 16 cores of an Intel Xeon Gold 6140 2.30GHz CPU with access to 32 GB of RAM. All experiments relied on version 1.4.2 of `MendelIHT` and Julia v1.5.4. IHT’s sparsity level *k* is tuned by cross-validation. The number of cross-validation paths is an important determinant of both computation time and accuracy. Thus, for simulated data, we employed an initial grid search involving 5-fold cross validation over the sparsity levels *k* ∈ {5, 10,…, 50}. This was followed by 5-fold cross-validation for *k* ϵ {*k*_*best*_ − 4, …, *k*_*best*_ + 4}. This strategy first searches the space of potential values broadly, then zooms in on the most promising candidate sparsity level. `GEMMA` and `mv-PLINK` were run under their default settings. For both programs, we declared SNPs significant w hose p-values w ere l ower than 0.05*/*24523. For `GEMMA`, we used the Wald test statistic.

### 3.5 Quality Control for UK Biobank

Using `MendelIHT.jl` we conducted two separate analyses on the second release of the UK Biobank [39], which contains ~ 500, 000 samples and ~ 800, 000 SNPs. Our first analysis deals with 3 hypertension traits: average systolic blood pressure (SBP), average diastolic blood pressure (DBP), and body mass index (BMI). Our second analysis deals with 18 NMR metabolomic quantitative traits related to total lipid levels (Figure 1). The featured metabolomic traits are available under category 220; their field IDs appear in Supplementary Table 7.

All traits were first log-transformed to minimize the impact of outliers. Then each trait was standardized to mean 0 and variance 1, so that the traits were treated similarly in mean-squared error (MSE) cross-validation. Following [8, 14, 19], we first filtered samples exhibiting sex discordance, high heterozygosity, or high SNP missingness. We then excluded samples of non-European ancestry and first and second-degree relatives based on empirical kinship coefficients. For 3-trait hypertension analysis, we also excluded samples who were on hypertension medicine at baseline. Finally, we excluded samples with < 98% genotyping success rate and SNPs with < 99% genotyping success rate and imputed the remaining missing genotypes by the corresponding sample-mean genotypes. Note that imputation occurs in IHT on-the-fly.

The final dataset contains 470, 228 SNPs and 185, 656 samples for the 3 hypertension traits and 104, 264 samples for the metabolomics traits. Given these reduced data and ignoring the Biobank’s precomputed principal components, we computed afresh the top 10 principal components via `FlashPCA2` [1] for the 3-trait analysis and `ProPCA` [2] for the 18-trait analysis. These principal components serve as predictors to adjust for hidden ancestry. We also designated sex, age, and age^2^ as non-genetic predictors.

## 4 Results

### 4.1 Simulation Experiments

Table 1 summarizes the various experiments conducted on the simulated data. For IHT, 5-fold cross-validation times are included. Multivariate IHT is the fastest method across the board, and is the only method that can analyze more than 50 traits. Multivariate IHT’s runtime increases roughly linearly with the number of traits. All methods perform similarly in recovering the pleiotropic and independent SNPs. Univariate IHT exhibits slightly worse true positive rate compared to multivariate methods. Given the identically distributed effect sizes in our simulations, all methods are better at finding pleiotropic SNPs than independent SNPs.

Notably, the false discovery rates (FDR) for both univariate and multivariate IHT are much lower than competing methods. Presumably, many of the false positives from mvLMM and CCA represent SNPs in significant LD with the causal SNP. IHT is better at distilling the true signal within these LD blocks because it considers the effect of all SNPs jointly. `GEMMA`’s mvLMM is better at controlling false positives than `mv-PLINK`’s CCA, but model fitting for mvLMM is slower, especially for large numbers of traits. In summary, IHT offers better model selection than these competitors with better computational speed.

### 4.2 3-trait UK Biobank Analysis

With 3 hypertension traits, the UK Biobank analysis completed in 20 hours and 8 minutes on 36 cores of an Intel Xeon Gold 6140 2.30GHz CPU with access to 180 GB of RAM. As described in the methods section, the featured traits are body mass index (BMI), average systolic blood pressure (SBP), and average diastolic blood pressure (DBP). A first pass with 3-fold cross-validation across model sizes *k* ϵ {100, 200,…, 1000} showed that *k* = 200 minimizes the MSE (mean squared error). A second pass with 3 fold cross-validation across model sizes *k* ϵ {110, 120,…, 290} showed that *k* = 190 minimizes the MSE. A third 3-fold cross-validation pass across *k* ∈ {191, 192,…, 209} identified *k* = 197 as the best sparsity level. Given *k* = 197, we ran multivariate IHT on the full data to estimate effect sizes, correlation among traits, and proportion of phenotypic variance explained by the genotypes.

IHT selected 13 pleiotropic SNPs and 171 independent SNPs. Selected SNPs and non-genetic predictors appear in the supplement as Tables 2-6. To compare against previous studies, we used the `R` package `gwasrapidd` [28] to search the NHGRI-EBI GWAS catalog [26] for previously associated SNPs within 1 Mb of each IHT discovered SNP. After matching, all 13 pleiotropic SNPs and 158 independent SNPs are either previously associated or are within 1Mb of a previously associated SNP. We discovered 3 new associations with SBP and 10 new associations associated with DBP. Seven SNPs, rs2307111, rs6902725, rs11977526, rs2071518, rs11222084, rs365990, and rs77870048, are associated with two traits in opposite directions.

**Table 2:** 13 pleiotropic SNPs selected by IHT listed with their effect sizes and sorted by their position on the chromosomes. An effect size of 0 means the particular predictor was not selected. The field *prior reports* records the number of SNPs previously associated with BMI, SBP, or DBP (p value < 10^−8^) that are within 1Mb of the given SNP. BMI = body mass index; SBP = systolic blood pressure; DBP = diastolic blood pressure.

| SNP | Chrom | BMI | SBP | DBP | # prior report |
| --- | --- | --- | --- | --- | --- |
| rs1801131 | 1 | 0.0 | 0.009 | 0.008 | 41 |
| rs17367504 | 1 | 0.0 | 0.02 | 0.019 | 41 |
| rs16998073 | 4 | 0.0 | -0.023 | -0.023 | 24 |
| rs1173727 | 5 | 0.0 | 0.016 | 0.017 | 19 |
| rs2307111 | 5 | 0.013 | 0.0 | -0.01 | 34 |
| rs6902725 | 6 | 0.0 | -0.009 | 0.01 | 4 |
| rs11977526 | 7 | 0.0 | 0.013 | -0.011 | 6 |
| rs2071518 | 8 | 0.0 | -0.008 | 0.009 | 6 |
| rs11222084 | 11 | 0.0 | -0.011 | 0.009 | 9 |
| rs3184504 | 12 | 0.0 | -0.011 | -0.018 | 41 |
| rs365990 | 14 | 0.0 | 0.008 | -0.011 | 6 |
| rs7497304 | 15 | 0.0 | -0.019 | -0.018 | 14 |
| rs77870048 | 16 | 0.0 | -0.011 | 0.01 | 20 |

**Table 3:** Non-genetic predictors selected by IHT listed with their effect sizes. PC is short for principal component. BMI = body mass index, SBP = systolic blood pressure, and DBP = diastolic blood pressure.

| covariate | BMI | SBP | DBP |
| --- | --- | --- | --- |
| intercept | 0.0002 | -0.0 | 0.0003 |
| sex | 0.1165 | 0.1639 | 0.1808 |
| age | 0.1073 | -0.1395 | 0.6484 |
| age<sup>2</sup> | -0.0505 | 0.4581 | -0.6131 |
| PC1 | -0.0287 | -0.0066 | -0.0074 |
| PC2 | 0.0052 | -0.0077 | -0.0011 |
| PC3 | 0.0174 | 0.0043 | -0.0091 |
| PC4 | -0.0019 | -0.0055 | 0.0015 |
| PC5 | 0.0022 | 0.0137 | 0.0167 |
| PC6 | -0.0165 | -0.0067 | -0.0056 |
| PC7 | -0.003 | -0.0038 | -0.0078 |
| PC8 | -0.0014 | -0.0015 | -0.0014 |
| PC9 | 0.0039 | -0.0004 | 0.0017 |
| PC10 | 0.0068 | 0.0005 | 0.0001 |

**Table 4:** 38 SNPs associated with BMI independently of SBP and DBP listed with their effect sizes and sorted by their position on the chromosomes. The field *prior reports* records the number of GWAS Catalog associations with BMI (p value < 10^−8^) that are within 1Mb of the given SNP.

| SNP | Chrom | $\beta$ | # Prior reports | SNP | Chrom | $\beta$ | # Prior reports |
| --- | --- | --- | --- | --- | --- | --- | --- |
| rs2815757 | 1 | 0.019 | 14 | rs17207196 | 7 | 0.016 | 9 |
| rs543874 | 1 | -0.028 | 22 | rs925946 | 11 | -0.007 | 24 |
| rs2820312 | 1 | -0.014 | 9 | rs6265 | 11 | 0.012 | 24 |
| rs62106258 | 2 | 0.027 | 51 | rs2049045 | 11 | 0.007 | 24 |
| rs11127485 | 2 | 0.017 | 51 | rs10835211 | 11 | -0.011 | 24 |
| rs13393304 | 2 | 0.008 | 51 | rs7138803 | 12 | -0.007 | 9 |
| rs713586 | 2 | -0.026 | 26 | rs7132908 | 12 | -0.014 | 9 |
| rs9821675 | 3 | 0.007 | 20 | rs4776970 | 15 | 0.01 | 17 |
| rs1062633 | 3 | 0.012 | 20 | rs16951304 | 15 | 0.008 | 17 |
| rs957919 | 3 | -0.015 | 17 | rs2531995 | 16 | 0.016 | 17 |
| rs61587156 | 3 | 0.017 | 9 | rs72793809 | 16 | -0.015 | 17 |
| rs34811474 | 4 | 0.015 | 3 | rs4788190 | 16 | 0.019 | 16 |
| rs10938397 | 4 | -0.022 | 17 | rs4889490 | 16 | 0.014 | 20 |
| rs13107325 | 4 | -0.026 | 7 | rs1421085 | 16 | -0.05 | 56 |
| rs2112347 | 5 | 0.009 | 22 | rs17782313 | 18 | -0.011 | 47 |
| rs1422192 | 5 | -0.015 | 20 | rs10871777 | 18 | -0.015 | 47 |
| rs2744962 | 6 | -0.017 | 33 | rs17773430 | 18 | -0.014 | 45 |
| rs987237 | 6 | -0.012 | 37 | rs1800437 | 19 | 0.017 | 20 |
| rs9473932 | 6 | -0.009 | 37 | rs3810291 | 19 | 0.022 | 12 |

**Table 5:** 66 SNPs associated with SBP independently of BMI and DBP listed with their effect sizes and sorted by their position on the chromosomes. The field *prior reports* records the number of GWAS Catalog associations with SBP (p value < 10^−8^) that are within 1Mb of the given SNP. A novel SNP is not within 1Mb of any GWAS Catalog associations.

| SNP | Chrom | $\beta$ | # Prior reports | SNP | Chrom | $\beta$ | # Prior reports |
| --- | --- | --- | --- | --- | --- | --- | --- |
| rs3936009 | 1 | 0.009 | 7 | rs12673516 | 7 | 0.01 | 3 |
| rs1757915 | 1 | -0.009 | 4 | rs2282978 | 7 | 0.016 | 4 |
| rs6684353 | 1 | 0.011 | 2 | rs2392929 | 7 | -0.027 | 5 |
| rs12069946 | 1 | -0.009 | 2 | rs2978456 | 8 | -0.01 | 1 |
| rs2820441 | 1 | 0.009 | 1 | Affx-32837790 | 8 | -0.009 | 3 |
| rs1522484 | 2 | 0.016 | 3 | rs35758124 | 8 | -0.01 | 2 |
| rs9306894 | 2 | 0.009 | 4 | rs10757278 | 9 | -0.01 | 3 |
| rs55654088 | 2 | -0.01 | 4 | rs10986626 | 9 | -0.009 | 5 |
| rs13002573 | 2 | 0.012 | 13 | rs12258967 | 10 | 0.009 | 10 |
| rs560887 | 2 | 0.011 | 1 | rs1908339 | 10 | 0.01 | 4 |
| rs10497529 | 2 | 0.011 | 3 | rs11191064 | 10 | 0.009 | 6 |
| rs1052501 | 3 | 0.02 | 6 | rs11598702 | 10 | -0.011 | 17 |
| rs2498323 | 4 | -0.008 | 3 | rs4980389 | 11 | -0.009 | 14 |
| rs776590 | 4 | 0.008 | 2 | rs573455 | 11 | -0.014 | 6 |
| rs17084051 | 4 | -0.01 | 3 | rs10750441 | 11 | 0.01 | 3 |
| rs1229984 | 4 | 0.009 | 1 | rs10770612 | 12 | 0.012 | 9 |
| rs6842241 | 4 | -0.008 | 2 | rs2681492 | 12 | 0.011 | 21 |
| rs4690974 | 4 | -0.01 | 9 | rs12882307 | 14 | 0.009 | 1 |
| rs13116200 | 4 | -0.01 | 2 | rs4903064 | 14 | -0.01 | 5 |
| rs7715779 | 5 | 0.007 | 12 | rs956006 | 15 | 0.009 | 1 |
| rs1173771 | 5 | 0.005 | 12 | rs34862454 | 15 | -0.012 | 14 |
| rs1982192 | 5 | 0.009 | Novel | rs3803716 | 16 | 0.008 | 3 |
| rs2303720 | 5 | 0.011 | 6 | rs4888372 | 16 | 0.01 | 6 |
| rs11954193 | 5 | 0.01 | 13 | rs60675007 | 16 | 0.008 | 2 |
| rs12198986 | 6 | -0.008 | 1 | rs9889363 | 17 | -0.011 | 6 |
| rs9349379 | 6 | 0.012 | 5 | rs185478092 | 17 | -0.01 | Novel |
| rs385306 | 6 | 0.011 | 7 | rs3744760 | 17 | -0.01 | 9 |
| rs12191865 | 6 | -0.014 | 3 | rs17608766 | 17 | -0.013 | 3 |
| rs1012257 | 6 | -0.009 | 3 | rs9909933 | 17 | 0.009 | 7 |
| rs9689048 | 6 | 0.009 | Novel | rs35688424 | 17 | -0.011 | 8 |
| rs2221389 | 6 | -0.012 | 1 | rs67882421 | 18 | -0.012 | 4 |
| rs9505897 | 6 | -0.01 | 1 | rs12459507 | 19 | -0.009 | 6 |
| rs57301765 | 7 | -0.014 | 2 | rs34328549 | 19 | 0.007 | 8 |

**Table 6:** 67 SNPs associated with DBP independently of BMI and SBP listed with their effect sizes and sorted by their positions on the chromosomes. The field *prior reports* records the number of GWAS Catalog associations with DBP (p value < 10^−8^) that are within 1Mb of the given SNP. A novel SNP is not within 1Mb of any GWAS Catalog associations.

| SNP | Chrom | $\beta$ | # Prior reports | SNP | Chrom | $\beta$ | # Prior reports |
| --- | --- | --- | --- | --- | --- | --- | --- |
| rs61776719 | 1 | -0.015 | Novel | rs4754834 | 11 | -0.009 | 1 |
| rs12739904 | 1 | -0.009 | 1 | rs59317921 | 11 | 0.009 | 2 |
| rs3766090 | 1 | -0.01 | Novel | rs7975252 | 12 | -0.009 | 1 |
| rs61822997 | 1 | -0.009 | Novel | rs7973748 | 12 | 0.011 | 10 |
| rs665834 | 1 | 0.011 | 1 | rs12581906 | 12 | 0.009 | 10 |
| rs2275155 | 1 | -0.013 | 6 | rs17287293 | 12 | 0.01 | Novel |
| rs11690961 | 2 | -0.012 | 1 | rs74340001 | 12 | -0.01 | 1 |
| rs10199082 | 2 | -0.01 | 3 | rs653178 | 12 | -0.007 | 16 |
| rs2692893 | 2 | -0.012 | 5 | rs12875271 | 13 | 0.011 | 3 |
| rs17362588 | 2 | -0.014 | 3 | rs36033161 | 14 | -0.009 | Novel |
| rs12996836 | 2 | -0.011 | 3 | rs686861 | 15 | -0.013 | 1 |
| rs1863703 | 2 | 0.009 | 3 | rs11853359 | 15 | 0.011 | 2 |
| rs2624847 | 3 | -0.009 | 3 | rs1378942 | 15 | -0.012 | 10 |
| rs3617 | 3 | 0.008 | 5 | rs4886615 | 15 | -0.008 | 10 |
| rs9850919 | 3 | 0.009 | 5 | rs2277547 | 15 | 0.01 | 1 |
| rs871606 | 4 | -0.018 | 1 | rs7174546 | 15 | -0.009 | 2 |
| rs1826909 | 4 | -0.01 | Novel | rs67456613 | 16 | -0.01 | 2 |
| rs1047440 | 5 | -0.014 | 3 | rs72790195 | 16 | -0.008 | 1 |
| rs11949055 | 5 | 0.009 | 5 | rs11078485 | 17 | -0.009 | 2 |
| rs1233708 | 6 | 0.013 | 2 | rs72824497 | 17 | 0.009 | 1 |
| rs11154022 | 6 | 0.01 | 3 | rs768168 | 19 | -0.011 | Novel |
| rs12110693 | 6 | -0.012 | 3 | rs997669 | 19 | -0.01 | 2 |
| rs9376740 | 6 | -0.009 | Novel | rs755690 | 19 | -0.01 | 1 |
| rs58023137 | 6 | -0.011 | 1 | rs2876201 | 20 | -0.009 | 6 |
| rs194524 | 7 | -0.012 | Novel | rs652661 | 20 | 0.015 | 6 |
| rs13226502 | 7 | -0.014 | 2 | rs78309244 | 20 | 0.014 | 6 |
| rs2469997 | 8 | 0.01 | 3 | rs6046144 | 20 | 0.015 | 2 |
| rs507666 | 9 | 0.012 | 2 | rs76701589 | 20 | 0.009 | 3 |
| rs3812595 | 9 | -0.009 | 1 | rs2831969 | 21 | 0.009 | Novel |
| rs183348357 | 10 | -0.009 | 7 | rs9982601 | 21 | -0.009 | 1 |
| rs3739998 | 10 | 0.005 | 1 | rs2298336 | 21 | 0.01 | 1 |
| rs2505083 | 10 | 0.007 | 1 | rs112005532 | 21 | 0.012 | 1 |
| rs7125196 | 11 | 0.01 | 5 | rs73167017 | 22 | 0.011 | 1 |
| rs2304500 | 11 | -0.012 | 1 |  |  |  |  |

One can estimate the genotypic variance explained by the sparse model as 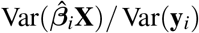 for each trait y_*i*_ where 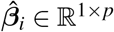 is the *i*th row of **B**. `MendelIHT.jl` outputs the values 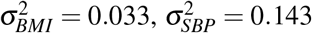, and 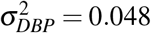. Note these estimates do not include contributions from the intercept or non-genetic predictors. The estimated correlations among traits are *r*_*BMI,SBP*_ = 0.197, *r*_*BMI,DBP*_ = 0.286, and *r*_*SBP,DBP*_ = 0.738. As expected, all traits are positively correlated, with a strong correlation between SBP and DBP and a weak correlation between BMI and both SBP and DBP.

### 4.3 18 trait UK Biobank analysis

A separate analysis of the 18 UK Biobank lipid traits finished in 53 hours on 32 cores of AMD EPYC 7502P 2.5GHz CPU with access to 252 GB of RAM. The peak RAM usage was 80.1 GB as measured by the seff command available on slurm clusters. Our cross validation search started with an initial grid of *k* ϵ {1000, 2000, 10000} and eventually terminated with *k* = 4678. The IHT run-time script with its detailed cross validation path is available in the Supplement.

Multivariate IHT found 218 independent and 699 pleiotropic SNPs for the 18 lipid traits. On average, a pleiotropic SNP is associated with 6.4 distinct lipid traits, suggesting that most significant SNPs for total lipid level are highly pleiotropic. Figure 1 depicts estimated effect sizes. The complete list of effect sizes as well as the estimated trait covariance matrix can be downloaded from our software page. The estimated proportion of variance explained for each trait 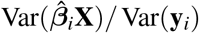 appears in the Supplement.

Although all traits are related to total lipids, we observe many associated genes contain distinct SNPs with opposite effects. Some of these reversals are caused by negatively correlated traits. Others are byproducts of IHT estimating the effect size of the alternate allele rather than that of the reference allele. Interestingly, SNP rs7679 has a large negative effect for Very Large HDL but a large positive effect for Small HDL despite the fact that the two traits are positively correlated. To verify this phenomenon, we conducted 18 univariate regressions considering only rs7679 plus an intercept. The result confirmed that this SNP indeed affects the two traits in opposite directions. SNPs such as rs7679 are compelling candidates for followup studies.

## 5 Discussion

This paper presents multivariate IHT for analyzing multiple correlated traits. In simulation studies, multivariate IHT exhibits similar true positive rates, significantly lower false positive rates, and better overall speed than linear mixed models and canonical correlation analysis. Computational time for multivariate IHT increases roughly linearly with the number of traits. Since IHT is a penalized regression method, the estimated effect size for each SNP is explicitly conditioned on other SNPs and non-genetic predictors. Analyzing three correlated UK Biobank traits with ~ 200, 000 samples and ~ 500, 000 SNPs took 20 hours on a single machine. A separate 18-trait analysis with ~ 100, 000 samples and ~ 500, 000 SNPs took 53 hours. IHT can output the correlation matrix and proportion of variance explained for component traits. `MendelIHT.jl` also automatically handles various input formats (binary PLINK, BGEN, and VCF files) by calling the relevant `OpenMendel` packages. If binary PLINK files are used, `MendelIHT.jl` avoids decompressing genotypes to full numeric matrices.

`MendelIHT.jl`’s superior speed is partly algorithmic and partly due to software/hardware optimization. Internally, each iteration of multivariate IHT requires a small *r* × *r* Cholesky factorization, where *r* is the number of traits. Each iteration also requires a dense matrix-matrix multiplication for computing gradients. For *r* ≤ 100 featured in this study, the factorization is trivial to compute. To speed up matrix multiplication, we developed a parallelized, tiled, and SIMD vectorized kernel that directly operates on binary PLINK files. This key innovation allows us to achieve performance near BLAS 3 calls without decompressing genotypes to numeric matrices. Because this kernel can be safely nested within IHT’s parallelized cross-validation step, we believe `MendelIHT.jl` is capable of utilizing hundreds of compute cores on a single machine.

IHT’s statistical and computational advantages come with limitations. For instance, it ignores hidden and explicit relatedness. IHT can exploit principal components to adjust for ancestry, but PCA alone is insufficient to account for small-scale family structure [32]. To overcome this limitation, close relatives can be excluded from a study. Additional simulations summarized in Supplemental Table 8 also suggest that analyzing traits of vastly different polygenic heritability may lead to slightly inflated false positive rates for the less polygenic traits. Thus, researchers may need to exercise caution when using mIHT for multiple traits when polygenic heritabilty differs by more than an order of magnitude. Although our simulation studies suggest the contrary, there is also the possibility that strong linkage disequilibrium may confuse IHT. Finally, it is unclear how IHT will respond to wrongly imputed markers and the rare variants generated by sequencing. In spite of these qualms, the evidence presented here is persuasive about IHT’s potential for multivariate GWAS.

We will continue to explore improvements to IHT. Extension to non-Gaussian traits is hindered by the lack of flexible multivariate distributions with non-Gaussian margins. Cross-validation remains computationally intensive in tuning the sparsity level *k*. Although our vectorized linear algebra routine partially overcomes many of the computational barriers, we feel that further gains are possible through GPU computing [21, 20, 22, 47]. In model selection, it may also be possible to control FDR better with statistical knockoff strategies [4, 36], especially if traits of vastly varying polygenicity are being considered. Given IHT’s advantages, we recommend it for general use with the understanding that genetic epidemiologists respect its limitations and complement its application with standard univariate statistical analysis.

## 6 Appendix

## 6.1 Significant SNPs from the UKB analysis

Tables 2-6 list the SNPs discovered by our 3-trait UK Biobank analysis. For our 18-trait analysis the selected SNPs are too numerous to list. However, all result can be accessed from our software page. Table 7 lists the proportion of variance explained for the 18 traits. The reported effect sizes correspond to the predictors of the log-transformed and standardized traits. To compare against previous studies, we searched the NHGRI-EBI GWAS catalog [26] using the `R` package `gwasrapidd` [28]. For each SNP discovered by IHT, we queried a 1Mb radius for other SNPs that have been previously associated with the given trait with p value < 10^−8^. Each known association is defined as the most significant SNP in a locus identified to be associated with the trait. All 13 pleiotropic SNPs were previously known and 158 out of 171 independent SNPs were previously known. Among the 13 newly discovered associations, 3 were with SBP and 10 were with DBP.

**Table 7:**
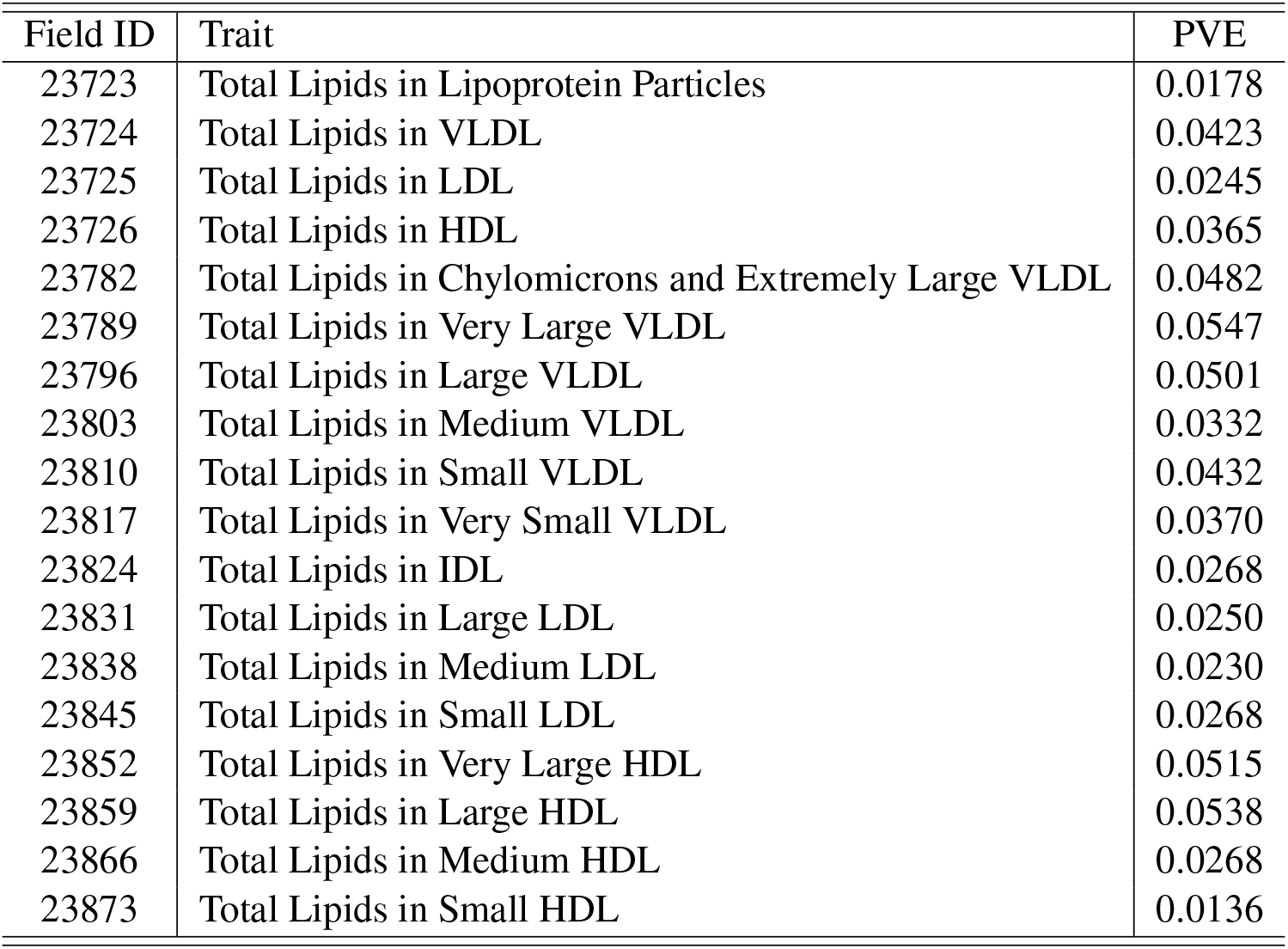
The estimated Proportion of phenotypic variance explained (PVE) for each of the 18 traits in the UK Biobank analysis.

| Field ID | Trait | PVE |
| --- | --- | --- |
| 23723 | Total Lipids in Lipoprotein Particles | 0.0178 |
| 23724 | Total Lipids in VLDL | 0.0423 |
| 23725 | Total Lipids in LDL | 0.0245 |
| 23726 | Total Lipids in HDL | 0.0365 |
| 23782 | Total Lipids in Chylomicrons and Extremely Large VLDL | 0.0482 |
| 23789 | Total Lipids in Very Large VLDL | 0.0547 |
| 23796 | Total Lipids in Large VLDL | 0.0501 |
| 23803 | Total Lipids in Medium VLDL | 0.0332 |
| 23810 | Total Lipids in Small VLDL | 0.0432 |
| 23817 | Total Lipids in Very Small VLDL | 0.0370 |
| 23824 | Total Lipids in IDL | 0.0268 |
| 23831 | Total Lipids in Large LDL | 0.0250 |
| 23838 | Total Lipids in Medium LDL | 0.0230 |
| 23845 | Total Lipids in Small LDL | 0.0268 |
| 23852 | Total Lipids in Very Large HDL | 0.0515 |
| 23859 | Total Lipids in Large HDL | 0.0538 |
| 23866 | Total Lipids in Medium HDL | 0.0268 |
| 23873 | Total Lipids in Small HDL | 0.0136 |

## 6.2 Additional simulation studies for IHT

In the main text, our simulation study explores the situation where causal SNPs are shared roughly equally among all traits. This is a reasonable assumption because we expect most multivariate GWAS to be conducted on similar traits, which are expected to share a similar number of causal variants. However, in practice, researchers may confront both highly polygenic traits along with very non-polygenic traits.

Here we design a simulation study with 3 traits, where each trait is perturbed by 10, 100, or 1000 causal SNPs, respectively. We use *n* = 10, 000 unrelated samples from the UK Biobank and restrict analysis to a genotype matrix constructed from 29481 SNPs on chromosome 10. Traits **Y**_*r*×10000_ are simulated just as in our main simulation study. We ignore the genetic relationship matrix since related samples have been filtered out. In summary,

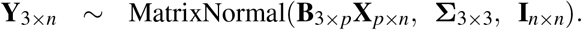

For the *r*th trait, *r* ϵ {1, 2, 3}, the effect sizes of the causal SNPs 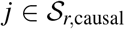 are Gaussian deviatres *β*_*j*_ ~ *N*(0, 0.1) with 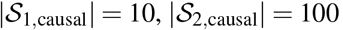, and 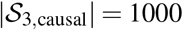. The causal SNP indices are chosen uniformly across the chromosome. The covariance between traits, Σ_3×3_, is generated as in our main simulation.

Table 8 reports the power and false discovery rate (FDR) for each trait separately, along with the overall power and FDR. Observe that FDR decreases as the number of causal SNPs *k* increases. Notably mIHT tends to select more SNPs than needed for a less polygenic trait if it is analyzed in unison with a highly polygenic trait. Obviously, this bias improves if we increase sample size or true effect sizes. Overall, these results suggest that one must be careful in analyzing multiple traits of vastly different polygenicity.

**Table 8:**
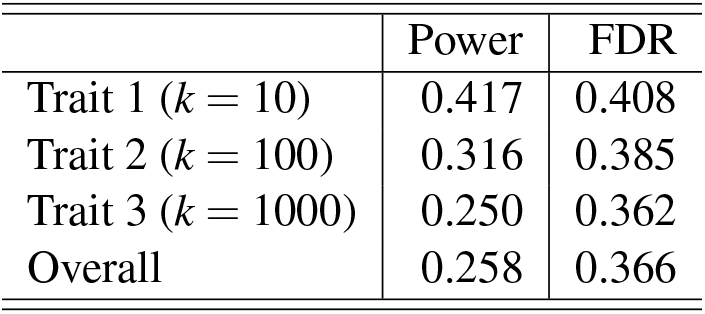
Multivariate IHT analyzing 3 simulated traits of different polygenicity. The 3 traits have 10, 100, and 1000 causal SNPs.

|  | Power | FDR |
| --- | --- | --- |
| Trait 1 ($k = 10$) | 0.417 | 0.408 |
| Trait 2 ($k = 100$) | 0.316 | 0.385 |
| Trait 3 ($k = 1000$) | 0.250 | 0.362 |
| Overall | 0.258 | 0.366 |

## 6.3 Loglikelihood

Consider multivariate linear regression with *r* traits under a Gaussian model. Up to a constant, the loglikelihood for the response vector y_*i*_ of subject *i* can be written

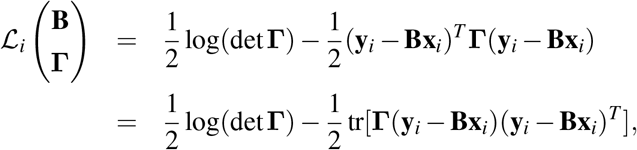

where **B** is the *r* × *p* matrix of regression coefficients, **x**_*i*_ is the *p* × 1 vector of predictors, and **Γ** is the *r* × *r* unstructured precision (inverse covariance) matrix. For *n* independent samples, let **Y** be the *r* × *n* matrix with *i*th column y_*i*_ and let **X** be the *p* × *n* design matrix with *i*th column **x**_*i*_. Then the loglikelihood for all samples is

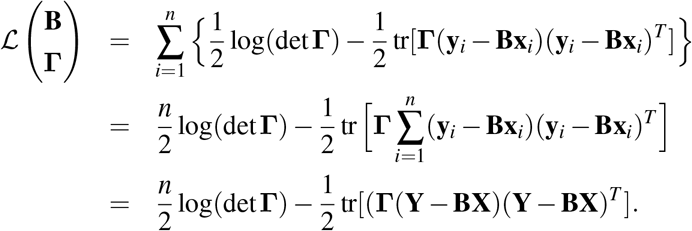

In subsequent sections we will present both full and block ascent IHT. The former updates **B** and **Γ** simultaneously. The latter alternates updates of **B** and **Γ**, holding the other parameter block fixed.

## 6.4 First Directional Derivative

Recall that the Hadamard’s semi-directional derivative [10, 23] of a function *f* (BOL) in the direction **v** is defined as the limit

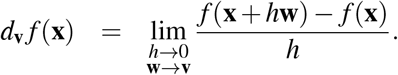

To calculate the directional derivative of the loglikelihood (3.1), we perturb **B** in the direction **U** and **Γ** in the symmetric direction **V**. The sum and product rules then give

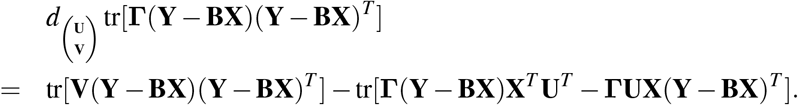

The directional derivative *d*_V_ lndet(**Γ**) = tr(**Γ**^−1^**V**) is derived in Example 3.2.6 of [23]. The trace properties tr(**CD**) = tr(**DC**) and tr(**C**^*T*^) = tr(**C**) consequently imply

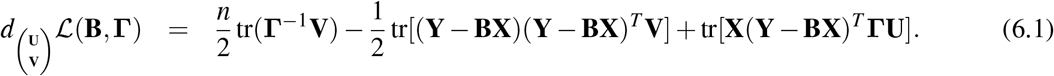

Because this last expression is linear in (**U, V**), the loglikelihood is continuously differentiable.

## 6.5 Second Directional Derivative

Now we take the directional derivative of the directional derivative (6.1) in the new directions 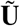 and 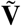. This action requires the inverse rule 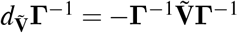 proved in Example 3.2.7 of [23]. Accordingly, we find

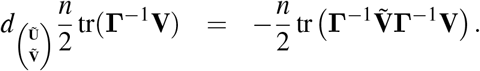

We also calculate

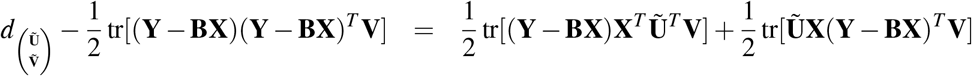

and

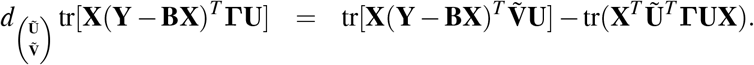

Finally, setting the two directions equal so that 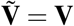 and 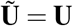 produces the quadratic form

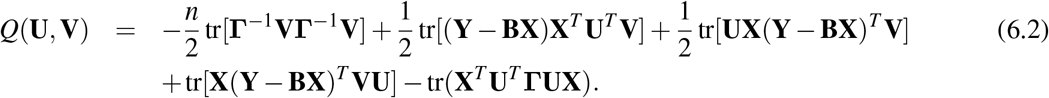

generated by the second differential.

## 6.6 Extraction of the Gradient and Expected Information

To extract the gradient from a directional derivative, we recall the identity *d*_v_ *f* (**x**) = ∇ *f* (**x**)^*T*^ **v** for vectors **v** and **x** and the identity tr(**A**^*T*^ **B**) = vec(**A**)^*T*^ vec(**B**) for matrices **A** and **B** [29]. The first identity shows that the directional derivative is the inner product of the gradient with respect to the direction **v**. The second displays the trace function as an inner product on dimensionally identical matrices. Thus, the matrix directional derivative is

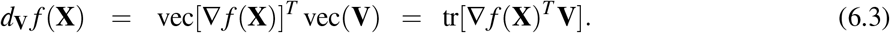

Inspection of the directional derivative (6.1) now leads to the gradient with blocks

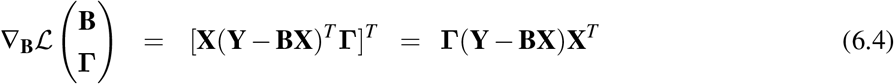

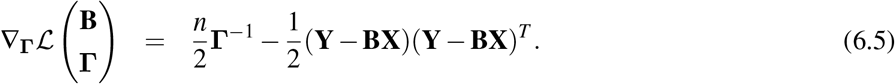

Analogously, the quadratic form (6.2) implicitly defines the Hessian **H** through the identity

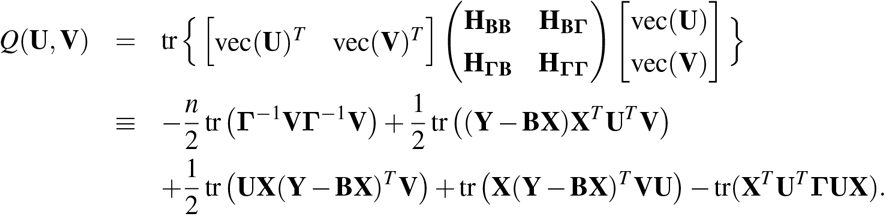

Because E(**Y**) = **BX**, the expected information **J** = E(−**H**) has the off-diagonal blocks 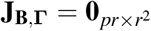 and 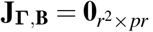 Now the Kronecker product identity vec(**ABC**) = (**C**^*T*^ ⊗ **A**) vec(**B**) implies

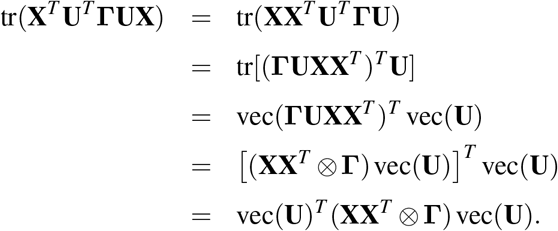

It follows that **J**_*BB*_ = **XX**^*T*^ ⊗ **Γ**. Similarly,

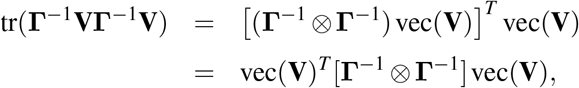

so that **J**_ΓΓ_ = **Γ**^−1^ ⊗ **Γ**^−1^. In summary, the expected information matrix takes the block diagonal form

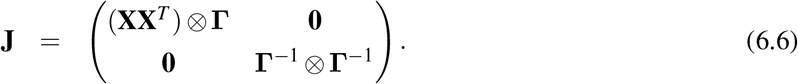

In our projected steepest ascent algorithm, the expected information matrix is never explicitly formed. It is implicitly accessed in the step-size calculation through the associated quadratic form *Q*(**B, Γ**).

## 6.7 Full IHT Step Size

The next iterate in full IHT is the projection of the point

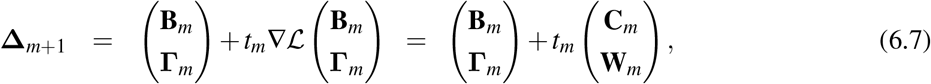

where 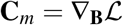 and 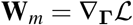 evaluated at (**B**_*m*_, **Γ**_*m*_). The loglikelihood along the ascent direction is a function of the scalar *t*_*m*_ and can be approximated by the second-order expansion

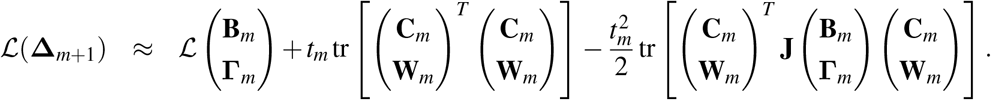

The choice

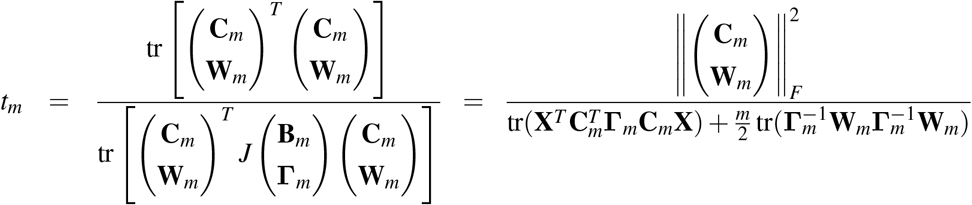

maximizes the approximation. If the support of the matrix (**B, Γ**) does not change under projection, then this IHT update is particularly apt.

## 6.8 IHT Projection

Recall that full IHT iterates according to

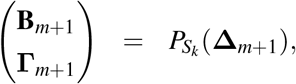

where Δ_*m*+1_ is derived in equation (6.7). Here *k* is a positive integer representing the sparsity level, which is assumed known. In practice *k* is found through cross-validation. The projection 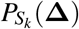 splits into separate projections for **B** and **Γ**. One can independently project each row of **B** to sparsity. Alternatively, one can require each row of **B** to have the same sparsity pattern if the same set of predictors plausibly contribute to all *r* traits. The **Γ** projection must preserve symmetry and positive semidefiniteness. Symmetry is automatic because the gradient of **Γ** is already symmetric. To project to positive semidefiniteness, one takes the SVD of **Γ** and project its eigenvalues *λ* to nonnegativity. One can even project **Γ** to the closest positive definite matrix with an acceptable condition number [40].

## 6.9 The Block Ascent IHT

In block ascent we alternate updates of **B** and **Γ**. The exact update (3.4) of **Γ** is particularly convenient, and we take advantage of it. Symmetry and positive semidefiniteness are automatically preserved. Inversion can be carried out via Cholesky factorization of **Γ**. This choice of **Γ** simplifies the step length

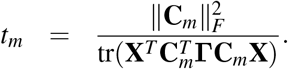

Note that the denominator of the step size does not require formation of the *n* × *n* matrix 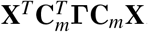. One can write 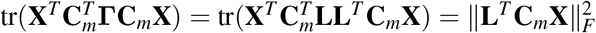, where **L** is the Cholesky factor of **Γ**. The matrix **L**^*T*^ **C**_*m*_**X** is fortunately only *r* × *n*.

## 6.10 UK Biobank Runtime Script

Here is the script used to perform our UK Biobank analysis

~~~
#
# Parameter explanations
# MvNormal: Distribution of traits is multivariate normal
# q: number of cross-validation folds
# min_iter: iterate at least 10 times before checking for convergence
#
~~~

~~~
using MendelIHT, Random, LinearAlgebra
BLAS.set_num_threads(1)
Random.seed!(2022)
plinkfile = “ukb.merged.metabolic.subset.european.400K.QC”
phenotypes = “traits.reordered.standardized.csv”
covariates = “covariates.reordered.standardized.csv”
~~~

~~~
# cross validate 1000, 2000, …, 10000
path = 1000:1000:10000
@time mses = cross_validate(plinkfile, MvNormal, path=path, q=3,
  covariates=covariates, phenotypes=phenotypes, min_iter=10,
  cv_summaryfile=“cviht.summary.roughpath1.txt”)
~~~

~~~
# cross validate 3100, 3200, …, 4900
k_rough_guess = path[argmin(mses)]
path = (k_rough_guess - 900):100:(k_rough_guess + 900)
@time mses = cross_validate(plinkfile, MvNormal, path=path, q=3,
  covariates=covariates, phenotypes=phenotypes, min_iter=10,
  cv_summaryfile=“cviht.summary.roughpath2.txt”)
~~~

~~~
# cross validate 4510, 4520, …, 4690
k_rough_guess = path[argmin(mses)]
path = (k_rough_guess - 90):10:(k_rough_guess + 90)
@time mses = cross_validate(plinkfile, MvNormal, path=path, q=3,
  covariates=covariates, phenotypes=phenotypes, min_iter=10,
  cv_summaryfile=“cviht.summary.roughpath3.txt”)
~~~

~~~
# cross validate 4671, 4672, …, 4689
k_rough_guess = path[argmin(mses)]
path = (k_rough_guess - 9):(k_rough_guess + 9)
@time mses = cross_validate(plinkfile, MvNormal, path=path, q=3,
  covariates=covariates, phenotypes=phenotypes, min_iter=10,
  cv_summaryfile=“cviht.summary.final.txt”)
~~~

~~~
# run full IHT on k = 4678
K = path[argmin(mses)]
@time iht_result = iht(plinkfile, K, MvNormal,
  summaryfile = “iht.final.summary.txt”,
  betafile = “iht.final.beta.txt”,
  covariancefile = “iht.final.cov.txt”,
  covariates=covariates, phenotypes=phenotypes, max_iter=2000)
~~~

## 7 Acknowledgements

This work is partially supported by National Institutes of Health grants T32-HG02536 (BC), R01-HG006139 (BC, KL, HZ), R35 GM141798 (KL, JS, HZ), R01-HG009120 (JS), K01DK106116 (JJZ), and R21HL150374 (JJZ); National Science Foundation grants DMS-1264153 (JS) and DMS-2054253 (HZ, JZ); and a National Research Foundation of Korea (NRF) grant 2020R1A6A3A03037675 (SK) from the Korean government (MSIT). The UCLA Institute for Digital Research and Education’s Research Technology Group supplied computational and storage services through its Hoffman2 Shared Cluster.

## 8 Author Contributions

KL conceived the project. BC, JS, and KL devised the methods and simulations. BC, SK, and HZ wrote the software. JZ and AJ accessed and analyzed the UK Biobank data. BC wrote the initial draft of the paper. All authors reviewed and edited the draft.

## 10 Web Resources

**Project name**: MendelIHT.jl

**Project home page**: https://github.com/OpenMendel/MendelIHT.jl

**Supported operating systems**: Mac OS, Linux, Windows

**Programming language**: Julia 1.6, 1.7

**License**: MIT

All commands needed to reproduce the following results are available at the `MendelIHT` site in the manuscript sub-folder. `SnpArrays.jl` is available at https://github.com/OpenMendel/SnpArrays.jl. `VCFTools.jl` is available at https://github.com/OpenMendel/VCFTools.jl. `BGEN.jl` is available at https://github.com/OpenMendel/BGEN.jl

